# The virulence and motility of *Pseudomonas syringae* pv. *actinidiae* mediated by a temperature sensor HktS-HktR

**DOI:** 10.1101/2024.11.20.624486

**Authors:** Yinan Xiao, Yifei Liang, Mingming Yang, Mingxia Zhou, Jing Huang, Nana Wang, Lili Huang

## Abstract

Kiwifruit bacterial canker (KBC), caused by *Pseudomonas syringae* pv. *actinidiae* (*Psa*) is one of the most devastating diseases of kiwifruit and can damage almost all kiwifruit varieties. The severity of the disease occurrence is closely related to the temperature. Our previous research indicated that *Psa* showed stronger pathogenicity and expansion ability at relatively cool temperatures, but how *Psa* senses environmental temperature and regulates its virulence mechanism remains unclear. In this study, 69 Histidine kinases (HK) in *Psa* were predicted through bioinformatics analysis, and 9 differentially expressed HK genes were identified at varying temperatures through pathogenicity detection and quantitative reverse transcription PCR (qRT-PCR). Among them, HktS as a temperature signal receiver was identified, and its response regulator (RR) HktR was determined through structure analysis and cotranscription assay. The results showed that HktR can bind to transcription factor RpoD, and RpoD bind to *hrpRS* promoter region, thus initiating the expression level of the type III secretion system (T3SS), which plays an important role in the pathogenesis of *Psa*. In addition, the motility of *Psa* was also regulated by HktS-HktR in a temperature-dependent manner. These findings reveal the molecular mechanism by which HktS-HktR acts as a temperature sensor to regulate bacterial virulence and motility of *Psa*, providing a new potential target for KBC control.

## Introduction

Different environmental conditions continuously stimulate bacterial growth and development, and temperature is a critical factor affecting bacteria. Therefore, bacteria must adjust their activities to adapt to temperature changes (Guijarro *et al*. 2015). Bacteria adjust gene expression patterns in response to changes in external temperature, causing them to undergo characteristic changes, and as a result, bacteria have developed precise and well-defined regulatory systems to regulate the expression of specific genes in response to changes in temperature (Guijarro *et al*. 2015; Schumann. 2012). Histidine kinases (HK) in bacteria can respond to changes in the extracellular environment (Wang *et al*. 2018). Signal transducing circuits mediated by HK play a central role in information processing. These circuits sometimes referred to as ‘two-component systems’ or ‘phosphorelays’; regulate a wide range of cellular responses, including bacterial chemotaxis, osmoregulation, photosensitivity, spore formation, plant responses to ethylene, and microbial pathogenesis (Bilwes *et al*. 1999). A two-component signaling system (TCS) typically consists of two or more multi-domain proteins, where HK is a homodimer, which utilizes ATP to transphosphorylate specific substrate histidine residues on adjacent subunits within the dimer (Yang and Inouye. 1991) and then transfers the phosphate groups on the histidine to specific aspartyl residues on the response-regulatory (RR) domain. HK can be divided into traditional orphan kinases and non-traditional mixed kinases. Orphan kinases have typically characteristic structural features, including a histidine residue capable of autophosphorylation and a phosphate group transfer domain. The structure of mixed kinases contains both HK and RR domains, enabling them to participate simultaneously in phosphorylation and dephosphorylation reactions (Sanders *et al*. 1992).

Currently, HK is described in *E. coli* as acting as a temperature sensor and regulating gene expression. Subsequent gene expression-related studies have linked the role of HK to cool-temperature gene regulation: temperature-related studies have been done in *Clostridium botulinum*, *Edwardsiella*, *Flavobacterium psychrophilum*, *Listeria monocytogenes*, plant pathogens *Agrobacterium*, *Antarctic Archaea*, as well as in *Pseudomonas syringae* and *Bacillus subtilis* (Najnin *et al*. 2016). In *Pseudomonas syringae*, a sensing HK (CorS) and two putative RR (CorR and CorP) were found. The transcription of genes for the synthesis of non-host-specific phytocoronary toxin (COR) biosynthesis was regulated through the interaction between CorS and its cognate RR CorR. Then, the virulence of *Pseudomonas syringae* was affected (Ullrich M. 1995).

Kiwifruit bacterial canker (KBC), caused by *Pseudomonas syringae* pv. *actinidiae* (*Psa*) poses a serious threat to the production of kiwifruit and can damage almost all kiwifruit varieties. The disease mainly affects the main trunk, branches and vines, new shoots and leaves, buds, and flowers of kiwifruit trees, which can cause the death of branches and vines in a short period, seriously restricting the healthy development of the kiwifruit industry, and it has become an important problem that needs to be solved urgently (Vinatzer *et al*. 2016). In the occurrence of KBC epidemics, temperature and rainfall factors are crucial in influencing the dynamics of pathogenesis (Beresford *et al*. 2017).

Temperature and humidity play an important role in the early and late onset of ulcer initiation, the rate of expansion, the degree of disease incidence, and the acceleration of disease spread, with cool temperatures and high humidity favoring the invasion and rapid multiplication of the pathogen. In Japan, *Psa* inoculated kiwifruit branches did not develop mottling at mean temperatures > 20°C (Serizawa *et al*. 1989), and using GFP-labelled *Psa*, *Psa* spread more rapidly in kiwifruit branches and leaf veins at cool temperatures (16°C) compared with 24°C (Vinatzer *et al*. 2016). However how *Psa* responds to temperature, regulates its growth, and pathogenicity is unknown.

The pathogenicity of *Psa* is dependent on its type III secretion system (T3SS), which produces the T3SS virulence effector, which the bacteria inject directly into plant host cells to interfere with host defense (Jin *et al*. 2003). The key regulator of T3SS in *Pseudomonas syringae* is HrpL, a member of the ECF alternative sigma factor family, which binds to highly conserved *hrp* box sequences within the promoters of T3SS-associated genes. Expression of *hrpL* is induced by HrpR and HrpS, two NtrC-like enhancer-binding proteins that bind to the *hrpL* promoter and recruit the sigma factor RpoN (σ^54^) and function as heterohexamers (Hutcheson *et al*. 2001; Jovanovic *et al*. 2011; Tang *et al*. 2006). This is known as the HrpR/S-HrpL-T3SS cascade reaction. T3SS gene expression in *Pseudomonas syringae* is coordinately regulated by a range of environmental and host signals (Tang *et al*. 2006), and T3SS gene expression is stimulated in cool osmotic pressure and acidic minimal media containing a carbon source (e.g., fructose), while it is repressed in most enriched media containing a complex nitrogen source or a broad spectrum of amino acid sources (Huynh. 1989; Xiao *et al*. 1992). Many studies have described the transcriptional regulation of the ‘HrpR/S-HrpL-T3SS’ cascade. Induction of the HrpRS manipulator is further regulated by the two-component system (TCS) RhpRS, CvsRS, and GacAS (Chatterjee *et al*. 2003; Fishman *et al*. 2018; Markel *et al*. 2016; Ortiz-Martín *et al*. 2010). These studies suggest that there may also be multiple regulators in *Psa* that regulate HrpRS transcription by recognizing different environmental or host signals, but how environmental signals are recognized and how HrpRS is regulated is not known.

In our previous study, we found significant differences in the growth, colonization, expansion ability, and pathogenicity of *Psa* at different temperatures (Wu *et al*. 2024), but it is not clear how *Psa* senses the environmental temperature and influences the virulence factors. In this study, we found that HktS-HktR can sense temperature and further initiate the transcription of *hrpRS* through binding to the downstream transcription factors RpoD In addition, HktS-HktR can also regulate bacterial swarming motility by sensing temperature. Overall, these findings not only confirm that HktS-HktR can sense temperature to regulate bacterial virulence but also play an important role in sensing temperature to regulate *Psa* swarming motility.

## Materials and methods

### Bacterial strains, plasmids, and growth conditions

Bacteria strains and plasmids used in this work are listed in Appendix A. The strongly pathogenic strain *Psa*_M228 was isolated from infected leaves of ‘HongYang’ in Shaanxi Province, China (Zhao *et al*. 2019). *Psa*_M228 was cultured at 28°C in Luria-Bertani (LB) medium. *Escherichia coli* (*E. coli*) strains were grown aerobically at 37°C in an LB medium. *E. coli* DH5α cells served as the host for constructing all recombinant vectors. *E. coli* BL21 (DE3) strain was used as the host to express recombinant proteins using pET30a and pGEX-6P- 1 vectors. Appropriate antibiotics were added to the culture as needed at the following concentrations: kanamycin (50 μgmL^-1^), gentamicin (25 μgmL^-1^), and ampicillin (100 μgmL^-1^). Electro-competent cells were prepared by Electroporation Buffer (Bio-Rad, USA). The transformation conditions of bacterial cells were set at 1.8 kV cm^-1^, 25 μF, and 200 Ω and conducted in a Bio-Rad Pulser XCell (Bio-Rad, USA) (Wang *et al*. 2019a).

### Bioinformatic analysis

The Pfam (Mistry *et al*. 2021) database contains the conserved domains of HATPase_c (Pfam02518) and Response reg (Pfam00072). The *Psa*_M228 proteome was scanned based on HATPase_c and Response reg using hmmsearch (Finn *et al*. 2011) on linux platform (parameter:-evalue 1e-10), and then the resulting proteins were built using hmmbuild to search for more mixed HKs. The newly created hmm model was used to perform a new round of hmmsearch on the *Psa*_M228 proteome (parameters:-evalue 1e-10) to obtain proteins as candidate mixed HK members. Candidate mixed HK proteins were analyzed by domain analysis using NCBI’s CDD search, and then the structural domains of each candidate protein were filtered: proteins containing both HATPase_c and Response reg structural domains were defined as mixed HKs and only HATPase_c was defined as an orphan HK. The structural domains of mixed HK proteins were mapped using the SMART database. The structural domains of mixed HK proteins were mapped using the SMART database. Based on the obtained mixed HK and orphan HK protein sequences, an evolutionary tree was constructed using MEGA (version 7.021, NJ method, bootstrap = 1000).

Protein sequence and DNA sequence of *Psa*_M228, and protein sequences of other strains were obtained from the National Centre for Biotechnology Information (NCBI) (https://www.ncbi.nlm.nih.gov/). These sequences were then submitted to NCBI-CDD (http://www.ncbi.nlm.nih.gov/Structure/cdd) and SMART (http://smart.embl-heidelberg.de/) to predict conserved functional domains (Letunic *et al*. 2015). Multiple alignments of protein sequences were performed by BLASTP (Thompson *et al*. 1994), CLUSTAL_W (Thompson et al. 1994), and ESprint 3.0 (Robert and Gouet. 2014).

### RT-PCR and qRT-PCR analysis

The relative expression levels of genes were detected by qRT-PCR, and Appendix B lists all the gene-specific primers used for qRT-PCR. Total bacterial RNA was extracted using the RNA pure Bacteria Kit (Jiangsu Cow in Biotech Co, Ltd, China). Briefly, 1 μg of total RNA was synthesized as a template for first-strand cDNA using the Revert Aid First Strand cDNA Synthesis Kit (Thermo Scientific, Massachusetts, USA), and then quantitative reverse transcription (qPCR) was performed using ChamQTM SYBR^®^ qPCR Master Mix (Vazyme, Q311, Nanjing, China). The internal control gene *gyrB* was used for normalization. The relative expression levels of three technical replicates were evaluated by the 2^-ΔΔCt^ method.

### Construction of insertion inactivation mutants

Deletion strains were constructed by an insertion inactivation method based on a homologous, one-step vector integration, using the suicide vector pK18mobSacB (Qian *et al*. 2008; Wang *et al*. 2013). Construction of the insertion inactivation mutant of 8 HK genes, *hktS*, *hktR*, and *rpoD* was based on the homologous, single cross-over method using suicide vector pK18mob. Genetic complementation of genes was performed by constructing recombinant vectors pBBR1-MCS5, which were provided in trans into corresponding bacterial strains. All primer sequences used in this study are listed in Appendix B.

### Phenotypic characterization of bacterial strains

Plant inoculation and virulence assays were conducted using two-year-old susceptible varieties of kiwifruit trees, *Actinidia chinensis* var. *chinensis* ‘HongYang’ as host plants, that were cultivated in a greenhouse at Northwest Agriculture and Forestry University, Shaanxi Province, China. Leaves were collected for pathogenicity determination. Plant inoculation and in planta bacterial growth assays were carried out according to previous reports (Zhao *et al*. 2019). Surface-sterilized healthy leaves were punched out of the leaf discs with a 10-mm-diameter punch and immersed in 30 mL of 1 × 10^5^ CFU · mL^-1^ bacterial suspension in a 50 mL tube. Vacuum penetration was carried out using a vacuum pump and a glass cover until the leaf disk sank to the bottom of the tube. Place 10 evenly infiltrated leaf discs into 0.1% water agar petri dishes. Healthy dormant kiwifruit woody canes without *Psa*_M228 were cut to 50 cm in length. After surface sterilization with 0.6% sodium hypochlorite for 10 min followed by 3 washes with sterile water, the woody canes were air-dried and carefully cut with a sterilized scalpel. In addition, 10 μL of a bacterial suspension at a concentration of 2 × 10^6^ CFU · mL^-1^ was inoculated into each wound area. Leaves and branches were incubated at 16°C and 30°C in a dark incubator. The phenotypes were observed and photographed at the 5 days post inoculation (dpi). The bacterial growth assay method was referred to a previously described procedure (Yuan and Xin. 2021).

### Swimming and swarming motility assays

The swimming and swarming motility of *Psa*_M228 and its mutant were determined on KB medium plates within semi-soft agar (0.25% and 0.4% agar), respectively (Río-Álvarez *et al*. 2013). The plates were prepared 1 h befer inoculation. Bacteria strains were cultured at 28°C overnight and adjusted to OD_600_ =1.0. The bacterial suspension (2.5 μL) was dropped on the center of the two types of KB plates, and incubated at 30°C and 16°C for 28 h, respectively. The experiment was performed three times, and each independent experiment involved three replicates.

### Model docking

The protein sequence was from NCBI, and the protein structure was predicted with ALPHAFOLD3 (Abramson *et al*. 2024). Molecular docking was performed with AutoDock Vina 1.2.0 and the docking results were analyzed by PLIP (Salentin *et al*. 2015) and visualized by PyMOL. The binding ability between the ligand and receptor was assessed by the Affinity value (kcal mol-1), with lower values indicating more stable binding.

### Protein expression and purification

Plasmids and primers used in this study are listed in Appendix B, which was used for cloning of His-tagged or GST-tagged proteins. The ORFs of proteins encoding genes were amplified from *Psa*_M228 genomic DNA and recombined into the pET30a plasmids. The recombinant plasmids were transformed into *E. coli* BL21 (DE3) strain and transferred into LB medium. Bacterial cells were induced at 16°C with 0.2 mM isopropyl β-D-1-thiogalactopyranoside (IPTG, Beyotime, Shanghai, China) for 10 h. The cultures were collected and suspended in 30 mL PBS (137 mM NaCl, 2.7 mM KCl, 10 mM Na_2_HPO_4_, 2 mM KH_2_PO_4_, pH 7.4, 1 mM PMSF). The bacterial pellets were lysed with sonication and centrifuged again at 12000 g for 30 min at 4°C to pellet bacterial debris. The supernatants were loaded into a Ni-NTA His•Bind® Resin column (Millipore Corp, USA), which had been equilibrated with PBS before use. The loaded Ni-NTA column was washed by 25 mL PBS. The target proteins were eluted with a gradient of 100 to 500 mM imidazole prepared in PBS.

For the purification of GST-tagged recombinant protein, the ORF was cloned from *Psa*_M228 genomic DNA into the pGEX-6P-1 vector. The resulting clone was transformed into the *E. coli* strain BL21 and induced at 28°C with 0.5 mM IPTG for 5 h. The bacterial pellet was sonicated in 30 mL PBS for 20 min at 4°C. The supernatant was collected by centrifugation at 12000 g for 30 min at 4°C and was incubated with the prepared GST•Bind Resin (Millipore Corp, USA) for 3 h at 4°C. Following incubation, untagged protein was The target proteins were eluted using elution buffer (20 mM reduced glutathione), and concentrated by centrifugation (Millipore, USA) at 4°C. The concentration of purified proteins was determined by the BCA protein assay kit (Solarbio, Beijing, China). Protein was assessed by sodium dodecyl sulfate-polyacrylamide gel electrophoresis (SDS-PAGE) and then stored at-80°C (Xie *et al*. 2021).

### Mass spectrometry analysis

Flag-tagged vectors are constructed and introduced into wild-type chromatin by transformation to extract total protein from the transformed species obtained. Add approximately 30 μL of anti-FLAG agarose (Abermat, Shanghai, China) to a 1 mL diluted sample of total protein (total protein/lysis buffer = 1:4) following the manufacturer’s instructions to capture the interacting proteins. The agarose was washed three times with 500 μL TBS (20 mM Tris-HCl, 500 mM NaCl, pH 7.5) and incubated overnight at 4°C. Proteins bound to the beads were eluted with 45 μL of elution buffer (0.2 M glycine, pH 2.5), and the eluent was immediately neutralized with 5 μL of neutralization buffer (1.5 M Tris, pH 9.0) (Qian *et al*. 2018). The eluted protein samples were identified by mass spectrometry at Beijing Protein Innovation Co., Ltd (Beijing, China).

### GST pull-down and Western blot analysis

Pull-down experiments involving a GST resin were performed as described earlier (Xu *et al*. 2018). The purified His-RpoD and GST-HktR proteins (5 μM each, final concentration) were added to 50 μL GST Resin (Beyotime, Shanghai, China) in 500 μL protein binding buffer (50 mM Tris-HCl, 200 mM NaCl, 1 mM EDTA, 1 mM DTT, 10 mM MgCl_2_, pH 8.0). For a negative control, GST was used in place of GST-HktR. The mixture tubes were incubated overnight at 4°C. The resin was collected by centrifugation at 12000 g for 1 min at 4°C and washed 6 times using protein binding buffer to remove non-specifically bound proteins. The proteins captured by GST Resin were eluted with Protein elution buffer (50 mM Tris-HCl, 400 mM NaCl, 1 mM EDTA, 1 mM DTT, 50 mM GSH, pH 8.0) and separated by SDS-PAGE.

For Western blots, the following antibodies were used: Goat Anti-Mouse (M21001S), His-Tag (M30111S), FLAG-tag monoclonal antibody (M20008S) (all from Abmart, Shanghai, China), GST-Tag (66001-2-Ig) (Proteintech, Wuhan, China).

### Microscale thermophoresis assay

The protein-DNA and protein-protein interaction were assayed by microscale thermophoresis (MST) using Monolith NT.115 (NanoTemper Technologies, Germany) as described earlier (Su *et al*. 2017; Xu *et al*. 2018). For the protein-protein binding assays, one of the proteins was labeled with the fluorescent dye NT-647-NHS (NanoTemper Technologies) via amine conjugation. Labeled proteins were incubated with unlabelled proteins to prepare test solutions. The test solutions were then drawn into capillary tubes (MO-K022) and measured in a Monolith NT.115. At least three biological replicates were performed for each assay and data were analyzed using MO. Affinity ANALYSIS v2.3.0 software.

### The Isothermal Titration Calorimetry

According to a previous study (Karonen, Oraviita, Mueller-Harvey, Salminen, & Green, 2015), isothermal thermal titration (ITC) was used to analyze the interaction between protein and DNA. Using the ultrafiltered protein, the amplified DNA is injected into the protein solution. Extensive degassing of the protein and DNA solutions was performed before injection into the cuvette and syringe. The ITC stirring speed was set to 300 rpm and ITC curves and thermodynamic data were analyzed at 25°C for various binding experiments. The collected data were analyzed using NanoAnalyzeTM software (TA Instruments, New Castle, USA) and binding isotherms were fitted using a binding model provided by the supplier.

### Electrophoretic mobility shift assay (EMSA)

To test the interaction between the *hrpRS* promoter region and its regulatory proteins, EMSAs were performed as described previously (Wang *et al*. 2019b). The spell-out FAM-labeled probes of the 60-bp promoter region of the *hrpRS* (Xu *et al*. 2018) were synthesized by Tsingke (Beijing, China). The DNA probe (30 ng) was mixed with protein in a 20 μL reaction mixture (5 mM Tris-HCl at pH 7.5, 1 mM MgCl_2_, 0.05 mM EDTA, 5 mM KCl, 0.3 μM HSA and 0.4% glycerol). After incubation for 30 min at 4°C, terminate the reaction with a 50% sucrose solution, the samples were loaded onto a 4% (w/v) non-denaturing acrylamide gel and subjected to electrophoresis at 100 V in 0.5 × Tris-borate-EDTA (TBE) buffer for 30 min at 4°C. EMSA signals (the labeled DNA fragments) were detected by Alex 488 nm using a VersaDoc imaging system (Bio-Rad, Philadelphia, USA).

## Results

### A *hktS* capable of sensing temperature identified from *Psa*

In order to screen HK capable of sensing ambient temperature, bioinformatics analysis was performed according to the genomic and proteomic sequences information of *Psa*_M228 HK family members of *Psa*_M228 were identified by scanning two-component signal transduction system-associated Pfam structural domains. Among them, proteins containing both HATPase_c and Response reg structural domains were defined as mixed HK, and those containing only HATPase_c were defined as orphan HK. Based on the protein sequences, the evolutionary tree was constructed using MEGA, and HK proteins were mapped according to the structural domains of proteins using the SMART database. The results showed that there are 69 HK in *Psa*_M228, including 21 mixed HK and 48 orphan HK genes (Fig. 1A, B).

**Fig. 1.**
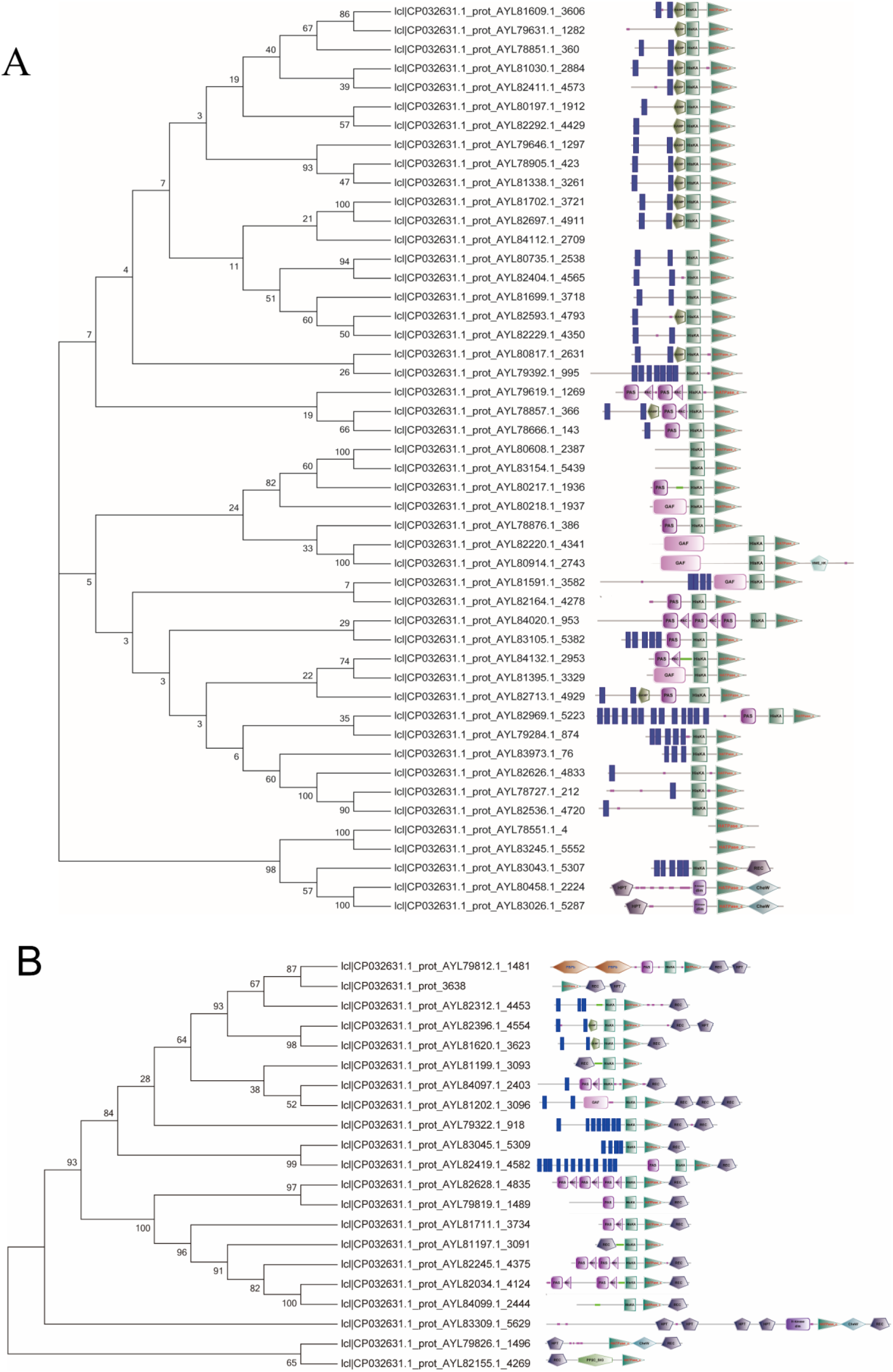
Histidine kinases predictive analysis in *Psa*_M228. Download HATPase_c (Pfam02518) and Response reg (Pfam00072) from the Pfam^1^ database. The conserved domains were individually scanned in the *Psa*_M228 proteome using hmmsearch^2^ on a Linux platform based on HATPase_c and Response reg. Subsequently, hmmbuild was used to construct the resulting proteins separately to identify more mixed HK. A new round of hmmsearch was then conducted on the *Psa*_M228 proteome using the newly created hmm model to search for proteins as candidate mixed HK members. Domain analysis of the candidate mixed HK proteins was performed using NCBI’s CDDsearch, along with the development of a Perl script to filter the domains of each candidate protein. Proteins containing both HATPase_c and Response reg domains were classified as mixed HK, while those containing only HATPase_c were classified as orphan HK. Based on the identified mixed HK, the sequences of HK proteins were utilized to build an evolutionary tree using MEGA (version 7.021, NJ method, bootstrap = 1000). The SMART database was leveraged to visualize the domains of the HK proteins. (A) Binding map of the mixed HK protein domain evolutionary tree. (B) Binding map of the orphan HK protein domain evolutionary tree.

The transcript levels of these HK genes in *Psa*_M228 were detected at 16°C and 30°C for 12 h. Nine HK genes expression was up-regulated significantly at 16°C compared to 30°C, including *psa_00715*, *psa_04875*, *psa_18850*, *psa_23100*, *psa_24500*, *psa_26820*, *psa_28560*, *psa_07600*, *psa_11300* (named as *hktS*) (Fig. 2A).

**Fig. 2.**
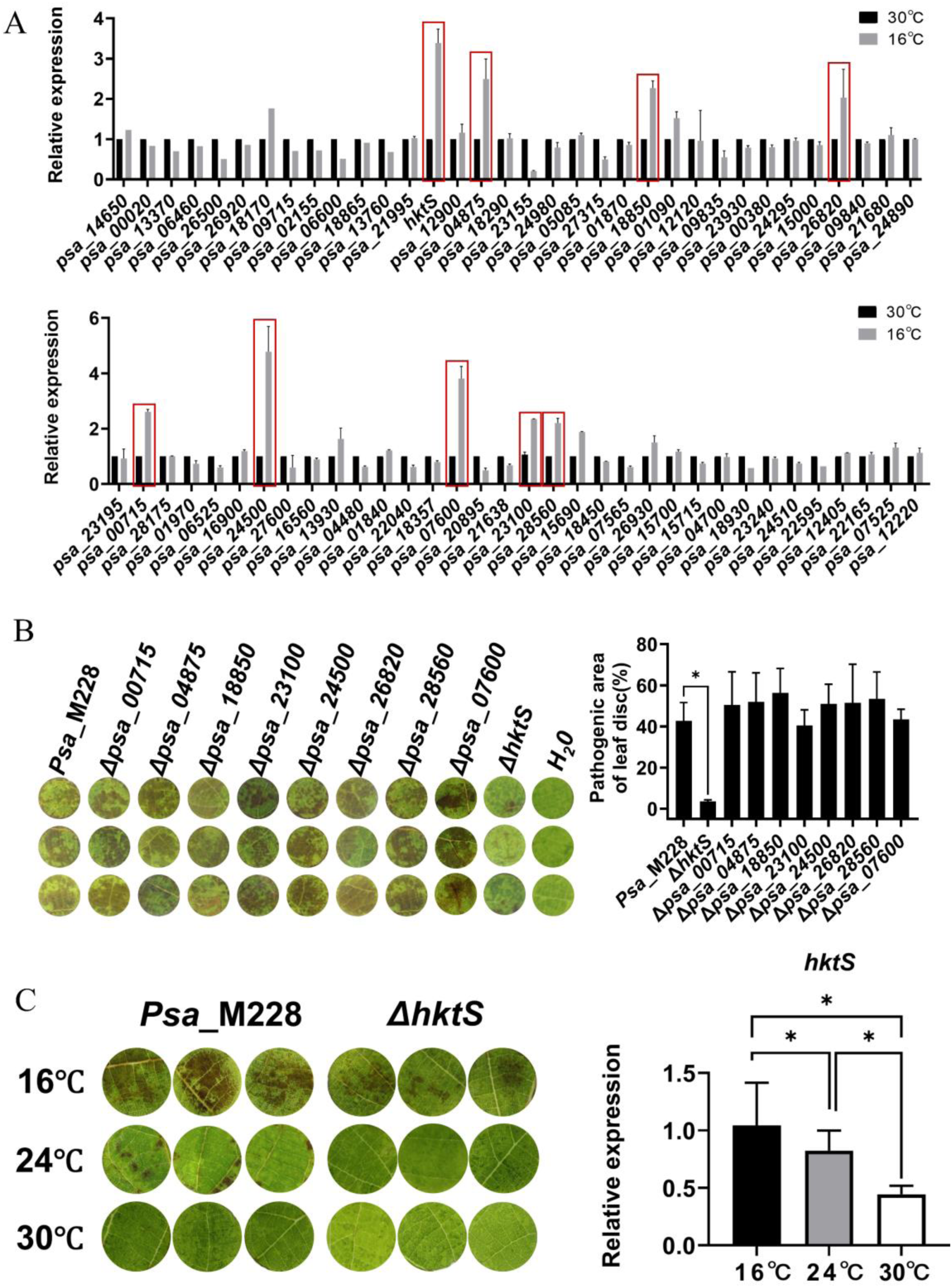
HktS selected as a temperature sensor from *Psa_*M228. (A) The expression of 68 HK genes of *Psa*_M228 at 12 h under different temperatures. (B) The pathogenicity of HK knockout mutant and wild-type strain *Psa*_M228 was tested by vacuum infiltration method, after 3 days of incubation, the lesion area was calculated; (C) The pathogenicity of strain *Psa*_M228 was tested by vacuum infiltration method at 16°C, 24°C and 30°C, respectively; The gene *htkS* expression levels in wild-type strain *Psa_*M228 are detected at 16°C, 24°C and 30 ° C, respectively. Asterisks: Student’s t-test difference was statistically significant (P < 0.05).

To determine whether these 9 HK can regulate the virulence of *Psa* in response to temperature, deletion mutants of the 9 HK genes were constructed in-frame (Appendix C), and their pathogenicity was tested. The results showed that the pathogenicity of mutant Δ*hktS* significantly reduced by comparison with the wild-type (WT) strain *Psa*_M228 (Fig. 2B). At the same time, the pathogenicity of *Psa*_M228 as well as the expression level of *hktS* in *Psa*_M228 changed significantly at different temperatures, however, these changes almost disappeared in the mutant Δ*hktS* (Fig. 2C). All the results indicated that *hktS* is a gene that can sense temperature and affect the pathogenicity of *Psa*.

### *hktS-hktR* sensing temperature regulates the virulence

HK must work in pairs with its downstream RR. The structure of HktS and its HktR (RR) were analyzed by the simplified modular architecture research tool (SMART, http://smart.embl-heidelberg.de/) online service. The HktS structure consists of a histidine phosphorylation transfer (HPT) structural domain, a signal-transducing histidine kinase homodimeric domain (H-kinase dim), an HK-type ATPase catalytic structural domain (HATPase_C), and a C-terminal CheW structural domain, HktR contains a receiver domain (REC) (Fig. 3A). The results of the cotranscription experiments showed that *hktS* and *hktR* have the same transcriptional orientation and belong to one operon, suggesting that they can be transcribed in pairs and exert functions (Fig. 3A).

**Fig. 3.**
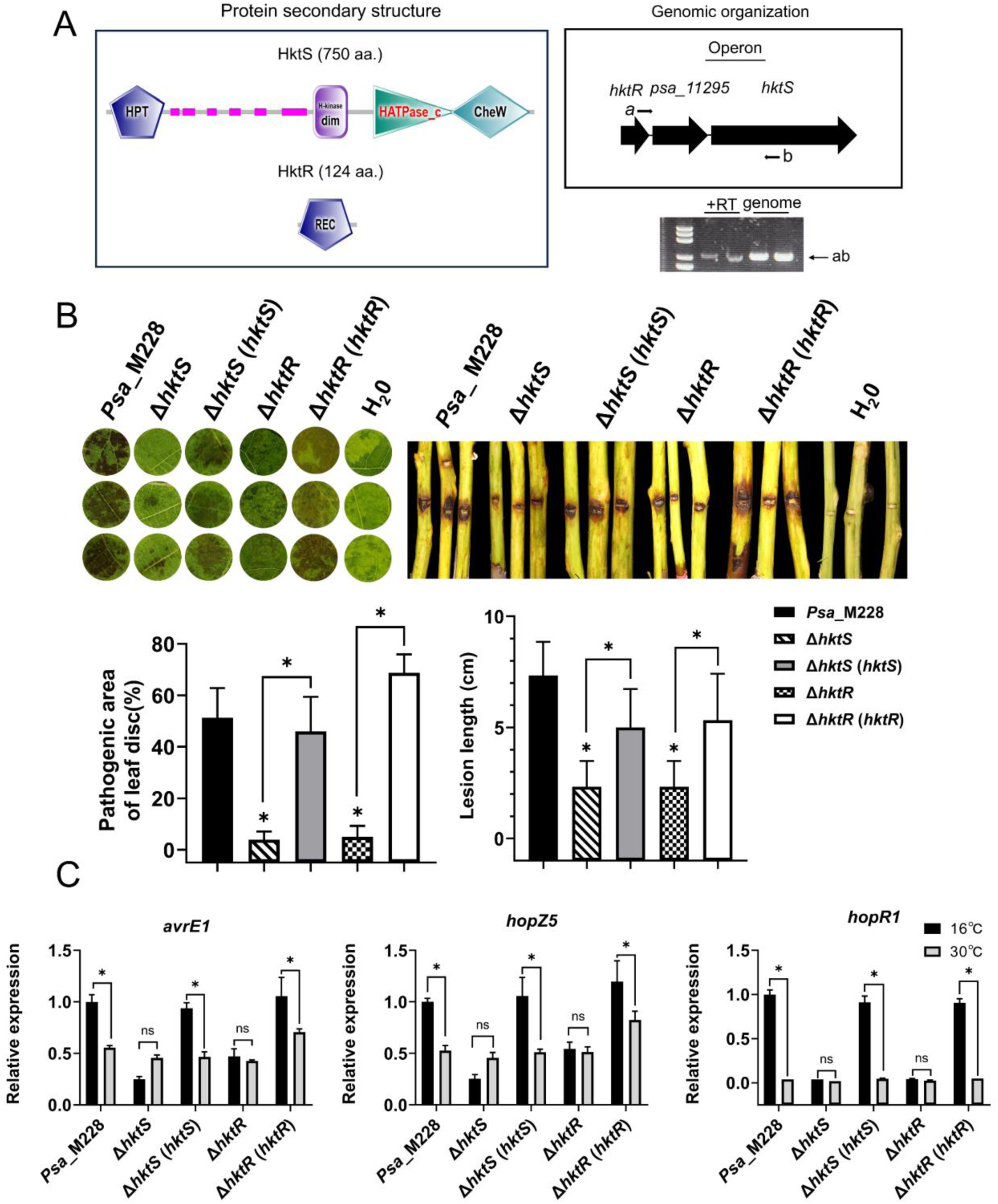
The virulence of *Psa*_M228 regulated by temperature sensor *hktS-hktR*. (A) Prediction of protein secondary structure by the SMART program. HPT: histidine phosphotransfer structural domain; H-kinase dim: signal transducing histidine kinase, homodimerisation structural domain; H-ATPase_C: ATPase catalytic structural domain; CheW structural domain: CheW-like domain.Schematic representation of the genomic organization of hktS-hktR: Large black arrows indicate genes and their transcriptional direction small arrows indicate the positions of primers used for RT-PCR; hktS and hktR constitute a bicistronic operon. +RT: cDNA as PCR template; genome: PCR template using the genome of Psa. The ab primers used to analyze RT-PCR for hktS-hktR are indicated by labeled arrows; (B) Vacuum infiltration and branch inoculation were used to test the pathogenicity of gene knock-out mutants and WT *Psa*_M228 strains. After incubation at 16°C for 3 days, the lesion area and length were measured. *Psa*_M228 was the wild-type strain and Δ*hktS* and Δ*hktR* were the deletion mutants Δ*hktS* (*hktS*) and Δ*hktR* (*hktR*) were the back-complemented strains; (C) The expression levels of T3SS genes (*avrE1*, *hopR1*, *hopZ5*) in WT and mutant strains at 16°C and 30°C. Asterisks: Student’s t-test difference was statistically significant (P < 0.05).

To reveal the effect of HktS-HktR on the virulence of *Psa_*M228, Deletion mutants of Δ*hktS*, Δ*hktR*, and complementary mutants Δ*hktS* (*hktS*), Δ*hktR* (*hktR*) were constructed (Appendix C). Due to the pathogenicity being most significant at 16°C, the pathogenicity test of all strains was determined at 16°C in the following experiments. In kiwifruit leaves and branches, WT and mutant strains were respectively inoculated. The pathogenicity of the Δ*hktS* mutant decreased by 92.14% and 68.19%, while that of the Δ*hktR* mutant decreased by 90% and 68.19%, and recovered in complemented mutants ΔhktS (*hktS*), Δ*hktR* (*hktR*) (Fig. 3B). In *Psa*_M228, the expression levels of the virulence-related T3SS genes (*avrE1*, *hopR1*, *hopZ5*) were downregulated by 2.5-fold, 2-fold, and 1-fold respectively, while the expression levels of the mutants Δ*hktS* and Δ*hktR* showed no significant difference at two different temperatures. The expression levels of the complemented mutants were restored to the WT state (Fig. 3C). Taken together, these findings suggest that *hktS-hktR* acts as positive regulators of bacterial virulence and are temperature dependent.

### *hktS-hktR* sensing temperature regulates the swarming motility

The motility ability of bacteria plays an important role in pathogen infection, colonization, and expansion, and is often influenced by environmental temperature. The swarming and swimming motility of *Psa*_M228 and mutants were determined at 16°C and 30°C. The swarming ability of *Psa*_M228 showed significant differences at two different temperatures, but no differences were observed in deletion mutants Δ*hktS*, Δ*hktR*. The effect of temperature on swarming ability has been restored in the complemented mutants Δ*hktS* (*hktS*) and Δ*hktR* (*hktR*). But temperature does not affect swimming ability (Fig. 4A).

**Fig. 4.**
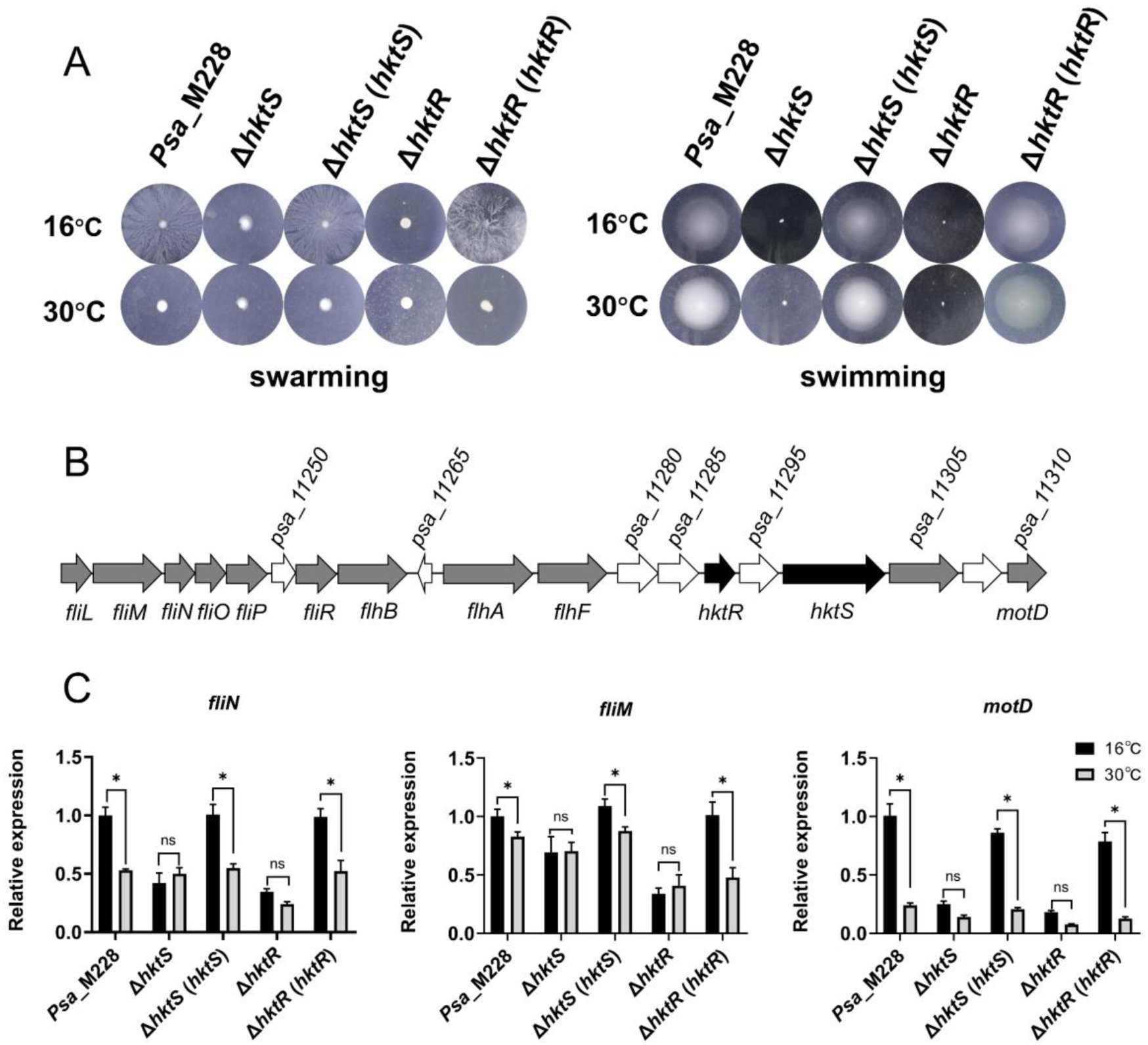
The swarming motility of *Psa*_M228 regulated by temperature sensor *hktS-hktR*. (A) Testing the swarming motility of mutant and WT strains at different temperatures in KB agar plates containing 0.45% agar. Testing the swimming motility of mutant and wild-type strains at different temperatures in KB agar plates containing 0.25% agar; (B) The *hktS-hktR* gene cluster shows the presence of many genes related to flagellar movement nearby; (C) *hktS-hktR* measures the expression levels of flagellar motor genes at different temperatures. Asterisks: Student’s t-test difference was statistically significant (P < 0.05).

To elucidate how *hktS-hktR* influences the group motility of *Psa_*M228, the expression levels of genes closely associated with bacterial swarming motility (*fliN*, *fliM*, *motD*) near *hktS-hktR* were examined at 16°C and 30°C (Fig. 4B). In *Psa*_M228, the expression levels of cluster movement-related genes were reduced by 2-fold, 1.25-fold, and 5-fold, while the expression levels of the mutants Δ*hktS* and Δ*hktR* showed no significant difference at the two temperatures. The expression levels of the complemented mutants were restored to the WT state (Fig. 4C). In summary, these findings indicate that *hktS-hktR* positively regulate bacterial swarming motility and are temperature-sensitive.

### HktR regulates the *hrpRS* promoter region through RpoD and influences T3SS expression

Since HktR lacks a detectable DNA-binding structural domain (Fig. 3A), we hypothesized that it controls the expression of T3SS genes through a transcription factor. To determine its downstream transcription factors, we conducted immunoprecipitation (IP) and mass spectrometry analysis, using HktR-FLAG as bait, which led to the identification of the transcription factor RpoD (Appendix D). It is a multi-structural domain subunit of bacterial RNA polymerase and is required for full virulence in bacteria. Based on the result, we hypothesize that HktR might bind to RpoD and then affect the pathogenicity of *Psa_*M228.

To verify the interaction between HktR and RpoD, we first performed molecular docking of HktR and RpoD using AlphaFold 3. The results showed that the amino acid residues Asn18, Arg14, Ser10, Arg87, Ala82, and Lys114 of HktR form hydrogen bonds with RpoD (Appendix E). Next, we expressed and purified HktR and RpoD (Appendix F). In GST pull-down experiments, purified HktR captured the RpoD protein RpoD-His_6_ (Fig. 5A). The dissociation equilibrium constant Kd of the protein interaction was determined to be 1.79 μM through MST experiments (Fig. 5B). Importantly, the GST tag itself did not bind to RpoD-His_6_ (Fig. 5A), and the BSA protein did not interact with fluorescently labeled GST-HktR (Fig. 5B).

**Fig. 5.**
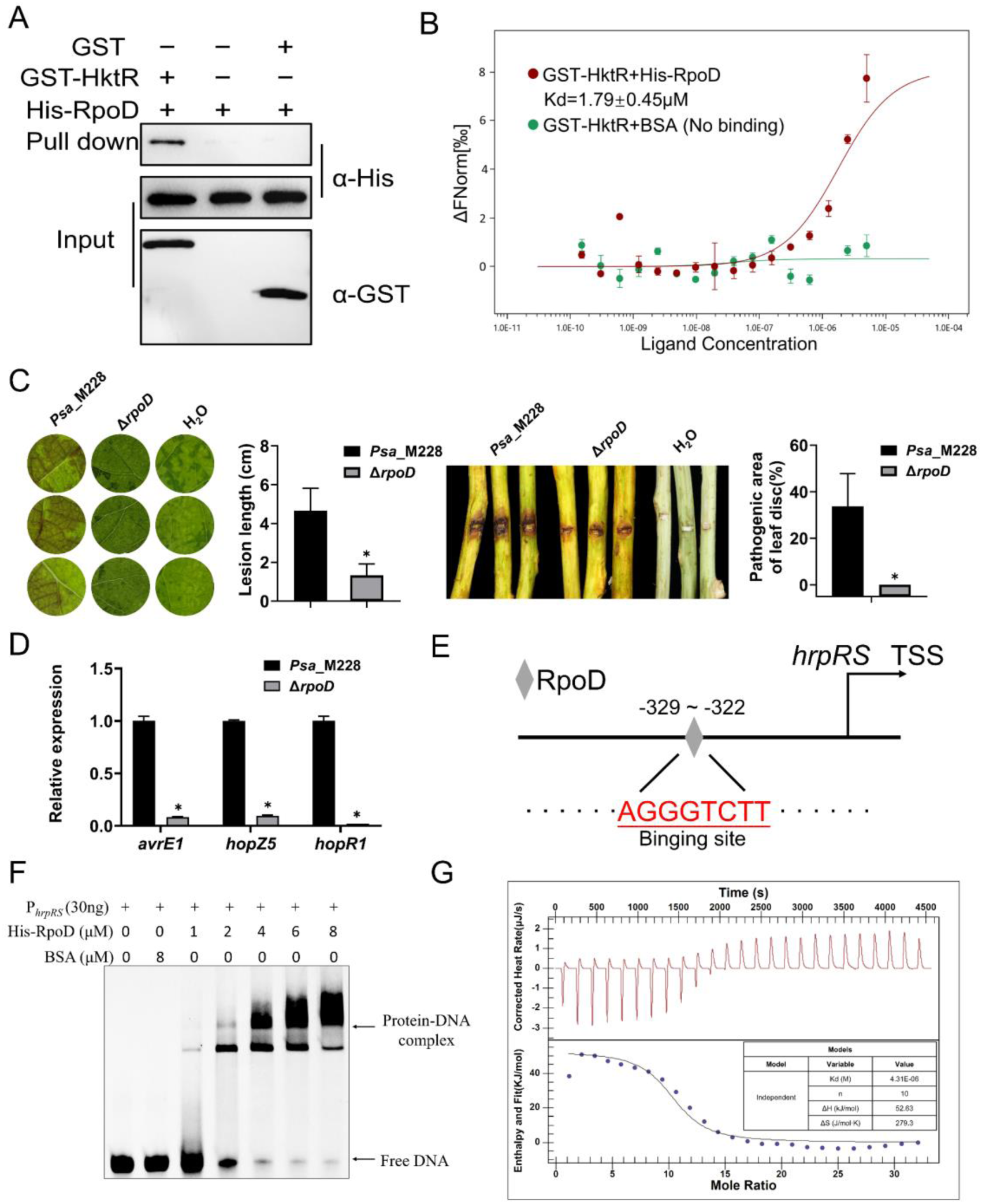
HktR regulates the *hrpRS* promoter region through RpoD and influences T3SS expression. (A) GST-HktR-RpoD-His pull-down assay confirmed that HktR interacts with RpoD, and GST was used as a negative control. western blot was performed using Anti-GST or Anti-His antibodies; (B) Through MST assay to analyze the affinity of HktR and RpoD. The purified GST-HktR is labeled with GST-Red dye, then co-incubated with purified His-RpoD in NT.115 standard capillary tubes for MST assay. BSA is used as a negative control. The binding curves and Kd values are calculated using MO.Affinity Analysis software; (C) Vacuum infiltration and branch inoculation were used to test the pathogenicity of *rpoD* knockout mutants and WT *Psa*_M228 strains. After incubation at 16°C for 3 days, the lesion area and length were measured. *Psa*_M228 represents the WT *Psa* strain, while Δ*rpoD* refers to the deletion mutants. (D) The expression levels of T3SS genes (*avrE1*, *hopR1*, *hopZ5*) in WT and mutant strains at 16°C and 30°C. Asterisks: Student’s t-test difference was statistically significant (P < 0.05). (E) Nucleotide sequence in the *hrpRS* promoter region. The predicted RpoD binding sites are colored and the putative RpoD binding sites are marked with red letters; (F) EMSA showed that RpoD binds to the *hrpRS* promoter in vitro. RpoD (1, 2, 4, 6, and 8 μM) was added to a reaction mixture containing 30 ng of probe DNA, and the reaction mixture was separated by polyacrylamide gel. Arrows indicate protein-DNA complexes and free DNA; (G) ITC showed RpoD binds to the *hrpRS* promoter in vitro. Use add 300ul of 18 μM Rpod to the titration cell, aspirate 2 mM HrpRS promoter region nucleic acid fragments and titrate.

To investigate the function of RpoD in *Psa*_M228, in-frame deletion mutants Δ*rpoD*, complementary mutants Δ*hktR* (*rpoD*) (overexpression of *rpoD* in mutant *hktR*) were constructed (Appendix C). Compared with *Psa_*M228, the pathogenicity of Δ*rpoD* did not exhibit little or no disease on host kiwifruit leaves and branches (Fig. 5C). The expression of the T3SS gene in Δ*rpoD* was down-regulated approximately 5-fold compared to the wild-type strain *Psa*_M228 (Fig. 5D). These results suggest that RpoD can positively regulate the expression of the bacterial virulence factor T3SS.

To investigate the relationship between HktR and RpoD, the T3SS gene expression levels were determined in *Psa*_M228, Δ*hktR*, and Δ*hktR* (*rpoD*) strain under different temperature conditions after 12 h of induction. The expression of the T3SS gene in the Δ*hktR* was significantly reduced and was restored in the Δ*hktR* (*rpoD*) compared with the WT strain *Psa*_M228. These findings indicate that HktR mediates the expression of the virulence factor T3SS through the downstream regulator RpoD (Appendix G).

To further determine how RpoD regulates T3SS gene expression, we analyzed the sequence of the RpoD protein in *Psa*_M228 as well as the sequences of the RpoD proteins of other bacteria. The BLASTP results showed that the M228_RpoD protein shares a highly conserved HTH structural domain with the RpoD proteins of other bacteria, such as in the *Pseudomonas* strain (Appendix H). There is a conserved RpoD-binding motif’AGGGTCTT’ in the *Psa*_M228 genome, which is located in the promoter region of *hrpRS* (Fig. 5E). Then the promoter region of *hrpRS was* cloned (Appendix I). EMSA and ITC assay both showed that *hrpRS* can bind to RpoD (Fig. 5F, G). These results suggest that RpoD can regulate T3SS by directly regulating the initiation of *hrpRS*.

## Discussion

Temperature has recently been shown to affect the development and virulence of a wide range of microbial species (Guijarro *et al*. 2015). Regulating the expression of various genes at the transcriptional level in response to temperature stimuli may be a way for microbial species to alleviate resource constraints and sustain life. HK, as a widely studied system that can sense external environmental signals, is widely distributed in bacteria, which accounts for 57.1% of the signal transduction proteins produced by *Pseudomonas syringae*, HK not only senses the external environment but most of the reported HK are also involved in T3SS, motility, and EPS production (Xie *et al*. 2022). Previous studies have found that CvsSR senses Ca^2+^signalling to regulate bacterial virulence and motility, RhpRS senses polyphenols to regulate bacterial virulence and metabolism, and VgrRS senses Fe^3+^, and that transcription of TdvA, the iron transport complex receptor, is inhibited by phosphorylated vgrR, which is required for the uptake of extracellular iron, in the presence of iron deficiency or a host plant environment (Desveaux *et al*. 2016; Fishman *et al*. 2018; Xie *et al*. 2022). As for temperature-sensing HK, including CorS in *Pseudomonas syringae* whose autophosphorylation is thought to occur via a temperature-dependent conformational change that results in the isolation of the catalytic cytoplasmic region containing conserved histidine into the membrane, thus acting as a thermosensitive switch (Braun *et al*. 2007); DesK in *Bacillus subtilis* senses the transfer of a phosphate group to the DesR and thus regulates the fatty acid desaturase gene DesK in *Bacillus subtilis* senses the transfer of a phosphate group to DesR to regulate the fatty acid desaturase gene, thereby controlling lipid saturation and maintaining membrane fluidity at cool temperatures, DesK can sense temperature by attaching to the cell membrane via the TMD (Albanesi *et al*. 2009); and LtrK and LtrR in the *Methanococcoides burtonii* sense temperature-regulated phosphotransfer (Najnin *et al*. 2016). However, these studies in HK have been at the structural level, and there is a paucity of information on how TCS senses temperature and then activates the expression of downstream virulence factors through a series of changes.

Our research data indicates that HktS-HktR is found to be compromised in bacterial virulence, particularly in bacterial secretion systems like T3SS, highlighting the direct involvement of the HktS-HktR system in regulating virulence genes in pathogens, playing a significant role in the pathogenic mechanism. The pathogenic mechanism mediated by HktS-HktR is closely related to temperature, as the loss of HktS-HktR eliminates the temperature influence on bacterial virulence (Fig. 3C), implying that HktR-HktS is a crucial temperature sensor that simultaneously affects the virulence of *P. aeruginosa*. However, the mechanism through which HktR-HktS transmits temperature signals is still unknown. Here, we propose a reasonable hypothesis that HktR may sense external temperature signals, stimulate autophosphorylation, and then transmit the phosphorylation signal to HktS. This assumption is based on the fact that most reported TCS perceive and transmit signals through phosphorylation transfer (Xie *et al*. 2022).

The motility of bacteria allows them to actively explore environments conducive to growth, thereby promoting proliferation. Movement typically involves invasion, migration, and disease development within the host (Abe *et al*. 2023). Therefore, understanding the impact of movement on virulence factors is crucial. Bacteria rely on flagella, type IV pili (TFP), and surfactants to coordinate movement on the surface of semi-solid culture media, a mode of movement referred to as swarming. Additionally, bacteria move through swimming facilitated by flagella in liquid environments (Zhang *et al*. 2022). In this study, we observed a significant reduction in both swarming and swimming motility after knocking out HktR (Fig. 4A). For the WT *Psa*_M228 bacteria, temperature primarily influences their swarming motility, leading us to speculate that HktS-HktR may regulate bacterial swarming motility by sensing temperature (Fig. 4A). Measurement of genes associated with swarming confirmed this hypothesis (Fig. 4C). Furthermore, studies suggest that bacterial swarming is linked to the overexpression of numerous virulence-related genes, including those encoding the T3SS and its effectors, extracellular proteases, and genes related to iron transport (Wang *et al*. 2019a). Based on these findings, we propose a reasonable hypothesis that upon temperature sensing, HktS-HktR may be involved in a mechanism related to regulating both bacterial swarming motility and virulence.

Recent evidence suggests that sigma (σ) factors can regulate bacterial virulence, σ factors play a key role in the recognition and binding of promoter DNA sequences by bacterial RNA polymerases during transcriptional initiation (Feklístov *et al*. 2014). The σ factors can be structurally and evolutionarily divided into two distinct families: σ^54^ and σ^70^ (Davis *et al*. 2017; Kazmierczak *et al*. 2005). The σ^70^ family consists of major sigma factors and alternative sigma factors that regulate the expression of genes involved in a variety of functions such as morphogenesis, iron uptake, and response to stressful environmental conditions (Bordes *et al*. 2011; Paget and Helmann. 2003). σ^70^ factor is also recognized as a transcription factor because of its role in activating and repressing gene expression at the transcriptional level in bacteria. The σ^70^ factor RpoD has been shown to mediate the T3SS by regulating *hrpG* and *hrpX* in *Xanthomonas oryzae*pv. *Oryzae* (Xu *et al*. 2023).

In this study, as a response regulator, HktR lacks the HTH structural domain, so it is speculated that it may bind to other proteins and thereby influence bacterial virulence. Screening the mass spectrometry data, several potentially interacting proteins with HktR were identified. Given that HktR can positively regulate bacterial virulence, considering the structural domains associated with HktR, it is hypothesized that it may interact with a transcription factor. Therefore, particular attention was paid to RpoD in the mass spectrometry data, and this interaction was further validated through techniques such as MST (Fig. 5B), revealing that HktR can interact with RpoD. Through measurements of pathogenicity and virulence factor expression, it was also confirmed that RpoD can positively regulate bacterial virulence (Fig. 5C, D). This discovery establishes a direct link between the two-component system HktS-HktR sensing temperature and the downstream transcription factor RpoD in the coordinated regulation of T3SS pathogenic mechanisms.

In *Xanthomonas oryzae*pv. *Oryzae* RpoD mediates the T3SS by regulating *hrpG* and *hrpX* (Xu *et al*. 2023), and it contains a conserved HTH structural domain that binds to the promoter regions of *hrpG* and *hrpX*. In the present study, we confirmed that the HTH structural domains of RpoD in *Psa* are highly conserved with those of RpoD in *Xanthomonas oryzae*pv. *oryzae* in which the HTH structural domain of RpoD is highly conserved (Appendix H), and a conserved motif located in the *hrpRS* promoter region for *rpod* binding was found in the *Psa*_M228 genome and demonstrated that RpoD can bind to this motif (Fig. 5F, G). In *Psa,*RpoD can regulate the T3SS via *hrpRS*. HktR binding to RpoD affects the T3SS.

According to our research findings, we found that HktS-HktR can act as a temperature sensor, regulating the motility and virulence of *Psa*_M228 (Fig. 6). HktS-HktR can sense temperature, modulating the expression of flagellar motor-related genes at the transcriptional level, thereby controlling bacterial swarming motility. Additionally, it can interact with the transcription factor RpoD and directly bind to the HrpRS promoter, inhibiting the downstream HrpRS-HrpL-T3SS pathway (Xie *et al*. 2019), which reduces the virulence of *Psa*. However, how HktS receives temperature signals and the interaction between HktS and HktR still need to be further investigated to improve this new temperature-responsive signaling pathway in *Psa*.

**Fig. 6.**
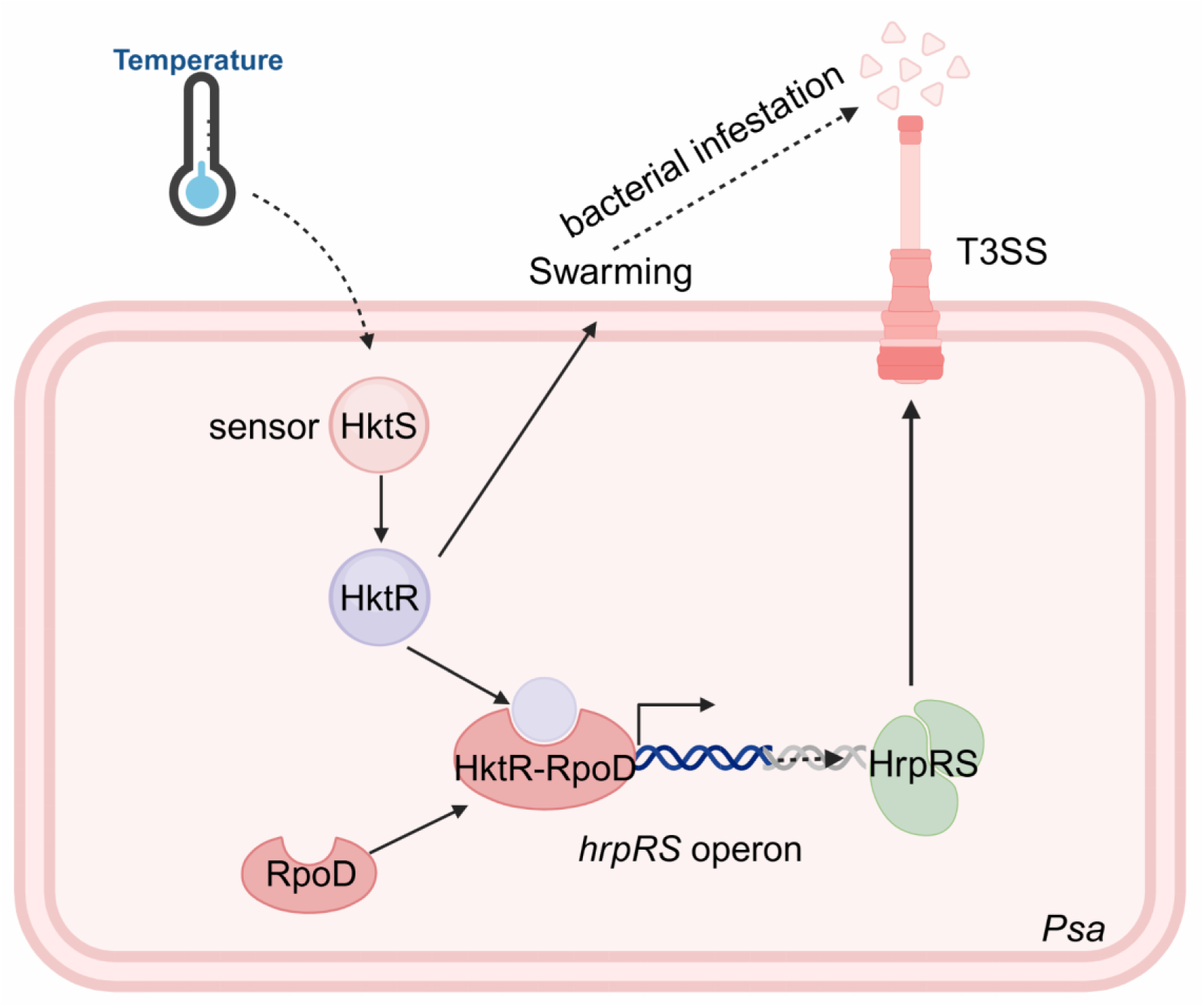
HktS-HktR, as a temperature sensor, mediates the regulatory model of motility and virulence in *Psa*_M228. The HktR-HktS two-component signalling system senses temperatures and subsequently initiates the transcription of the *hrpRS* gene by binding to the downstream transcription factor RpoD. Additionally, HktS-HktR can sense temperature to influence bacterial swarming motility, thereby affecting bacterial infection. In this study, we identified HktS-HktR as a temperature sensor that regulates gene expression in response to temperature stimuli in the kiwifruit bacterial rot disease *Psa*.

## Acknowledgements

This work was supported by the National Natural Science Foundation of China (32100148), National Key R&D Program of China (2022YFD1400200) and the Special Support Plan for High-level Talent of Shaanxi Province.

## Author contributions

LH and NW designed the research. YX, YL, MZ and JH performed the experiments. YX, NW and MY analyzed the data. YX and NW wrote the manuscript. All authors discussed the results and commented on the manuscript before submission.

